# Sleep problems are associated with hypersensitivity to touch in children with autism

**DOI:** 10.1101/255323

**Authors:** Tzischinsky Orna, Meiri Gal, Manelis Liora, Bar-Sinai Asif, Fluser Hagit, Michaelovski Analya, Zivan Orit, Ilan Michal, Faroy Michal, Menashe Idan, Dinstein Ilan

## Abstract

**Background:** Sensory abnormalities and sleep disturbances are highly prevalent in children with autism, but the potential relationship between these two domains has rarely been explored. Understanding such relationships is important for identifying children with autism who exhibit more homogeneous symptoms.

**Methods:** Here we examined this relationship using the Caregiver Sensory Profile and the children’s sleep habit questionnaires, which were completed by parents of 69 children with autism and 62 frequency age-matched controls.

**Results:** In line with previous studies, children with autism exhibited more severe sensory abnormalities and sleep disturbances than age-matched controls. The sleep disturbance scores were strongly associated with touch and oral sensitivities in the autism group and with touch and vestibular sensitivities in the control group. Hyper sensitivity towards touch, in particular, exhibited the strongest relationship with sleep disturbances in the autism group and single-handedly explained 24% of the variance in sleep disturbance scores. In contrast, sensitivity in other sensory domains such as vision and audition was not associated with sleep quality in either group.

**Conclusions:** While it is often assumed that sensitivities in all sensory domains are similarly associated with sleep problems, our results suggest that hyper sensitivity towards touch exhibits the strongest relationship to sleep disturbances when examining children autism. We speculate that hyper sensitivity towards touch interferes with sleep onset and maintenance in a considerable number of children with autism who exhibit severe sleep disturbances. Studies that examine the effects of tactile sensory therapies/aids on sleep quality and behavioral improvement in these children are, therefore, highly warranted.

## Background

Autism is a remarkably heterogeneous disorder where different individuals exhibit distinct behavioral symptoms. This heterogeneity is apparent in the core symptoms that define the disorder (i.e., impaired social communication/interaction, repetitive behaviors, and restricted interests) and in additional symptoms that are prevalent in individuals with autism (e.g., sensory abnormalities and sleep disturbances). A major goal of contemporary autism research is to identify individuals who share more homogeneous symptoms and who may benefit from targeted interventions [1,2]. Understanding potential relationships across symptom domains is an important step in characterizing individuals with more homogenous symptoms.

A large body of literature has shown that sensory problems are apparent in 60-90% of individuals with autism [3–8]. This has motivated the addition of sensory problems as a diagnostic criterion of autism in the DSM-5 [9]. However, sensory problems in autism can vary widely and include both hypo and hyper sensitivities in different sensory modalities [3,5,10–14]. The severity of sensory symptoms do not seem to correlate with cognitive abilities, adaptive behaviors, or autism severity [7]. Nevertheless, might the existence of particular sensory abnormalities in specific individuals with autism inform us about their other behavioral symptoms, such as sleep problems?

Sleep disturbances are also very common in individuals with autism with a prevalence of 40-80% [15–18]. Disturbances include difficulty falling asleep, frequent wakings during the night, shorter sleep duration, restlessness during sleep, and difficulty falling asleep. Previous studies have reported that sleep disturbances are more prevalent in regressive autism [17], increase with autism severity [16,19], and may [20,21] or may not [15,18] be associated with cognitive levels. In addition, the severity of sleep disturbances in children with autism seems to scale with their level of anxiety, attention deficits, impulsivity, challenging behaviors, and the use of medication [16,19,22–24]. Several studies have hypothesized that sensory sensitivities are also expected to correlate with sleep problems in autism [17,25], but this potential relationship has rarely been examined empirically.

With this in mind, two recent studies used Autism Speaks’ Autism Treatment Network (ATN) to examine the potential relationship between sensory abnormalities and sleep disturbances in autism. The ATN is a large national database [26] containing a wide variety of behavioral information from children with autism, which includes the Short Sensory Profile and the Child Sleep Habit Questionnaire (CSHQ). Both studies reported that children’s anxiety levels were significantly correlated with sleep disturbances. In addition, one study reported a significant relationship between the under-responsive/sensory-seeking or the auditory filtering subscale scores of the Short Sensory Profile and the total sleep disturbance scores of 23 item version of the CSHQ. While significant, both of the Short Sensory Profile predictors explained only 1% of the variance in the sleep disturbance scores [19]. The second study focused on quantifying sensory over-responsivity by summing the scores of questions in the Short Sensory Profile that pertain to sensory sensitivities in several domains (i.e. touch, vision, taste/smell, and audition). This study reported significant correlations between sensory over-responsivity and sleep-onset delay, sleep duration, and night awakenings, but not with the other CSHQ sub-scales [23].

The goal of the current study was to extend previous findings by examining data from the complete Caregiver Sensory Profile, which allow for more detailed analyses of sensory hypo and hyper sensitivities in each of five sensory domains (audition, vision, taste/smell, vestibular, and touch). We hypothesized that sleep disturbances, as assessed by the CSHQ, may be more strongly associated with sensory abnormalities in specific sensory domains rather than with Short Sensory Profile sub-scales that integrate scores across the sensory domains [27].

## Methods

### Participants

A total of 131 children participated in study: 69 children with autism (age: 3 to 7; mean age: 4.94±1.23; 56 male) and 62 frequency matched controls (mean age: 4.82±1.15; 41 males). There was no significant difference in the age of participating children across groups (t(129)) = 0.64, p = 0.57, two tailed t-test). Children with autism were recruited through the Preschool Psychiatry Unit and the Child Development Institute at Soroka Medical Center [28]. Control children of the same age were recruited from the community using an online forum at the university. Parents of all control children reported that their children were never suspected of having any developmental problems. The Helsinki committee at Soroka Medical Center approved this study and parents of all participating children signed an informed consent form.

#### Diagnosis

All children with autism met the DSM-5 criteria for autism as determined by both a physician (Child Psychiatrist or Neurologist) and a developmental psychologist [28]. 49 of the 69 children with autism also completed an Autism Diagnostic Observation Schedule (ADOS) assessment to confirm the diagnosis [29]. 22 of the 69 children with autism were taking medications that included Melatonin, Risperdal, Ritalin, and Neuleptil.

**Table 1:**
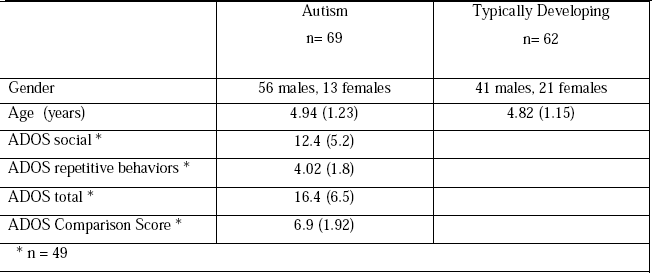
Sample characteristics. Gender and age of autism and control children as well as ADOS scores from the 49 children with autism who completed the assessment.

### Sensory Profile

We used the Hebrew version of the Caregiver Sensory Profile questionnaire to assess sensory sensitivities in all children [30]. This questionnaire contains 125 questions that quantify the frequency of abnormal behavioral responses to various sensory experiences [31]. In the current study we focused on questions that assess auditory, visual, vestibular, touch, and oral sensory processing. These questions are split into high threshold and low threshold items, which measure hypo and hyper sensitivities respectively, with lower scores indicating more severe symptoms. We examined differences across groups for each of these scores separately and also for the total raw score, which combines both low and high threshold items to indicate general sensory abnormalities while keeping in mind that some children exhibit both hypo and hyper sensitivities to different sensory experiences (Tables 2&3).

### Children’s Sleep Habits Questionnaire (CSHQ)

Parents of all participants scored their child’s sleep behaviors using the Hebrew version of the CSHQ [32,33]. This caregiver questionnaire includes 33 items which are divided into eight subscales representing different sleep disturbances: bedtime resistance, sleep anxiety, sleep-onset delay, sleep duration, night wakings, daytime sleepiness, sleep-disordered breathing, and parasomnias. The scores of these 8 domains are summed to generate a total score of sleep disturbances for each child. The internal consistency values of the CSHQ sub-scales in our study were α=0.533−0.758, and their consistency with the total sleep score was α=0.838.

### Statistical analyses

All statistical analyses were performed with Matlab (Mathworks, USA). We performed two-tailed t-tests with unequal variance to compare the 5 sensory measures (visual, auditory, vestibular, touch, and oral) of the Sensory Profile across the autism and control groups. Equivalent tests were performed for the 9 sleep measures (bedtime resistance, sleep onset delay, sleep duration, sleep anxiety, night wakings, parasomnias, sleep disordered breathing, daytime sleepiness, and total sleep score). All tests were corrected for multiple comparisons using the Bonferroni method (i.e., 5 comparisons in the Sensory Profile and 9 in the CSHQ). All sensory and sleep measures were close to normal distributions as demonstrated by their skewness and kurtosis values, which were all between −1 and 1, except for the visual and vestibular scores in the control group (skewness = −1.48 and −1.23, kurtosis = 4.38 and 1.59, respectively). Potential relationships between scores from each of the sensory modalities and the total sleep score were examined using both Pearson’s and Spearman’s correlations. This ensured that our conclusions were not based on the assumption that the distributions of the variables were normal or that the relationships were linear.

In a final set of analyses, we performed several regression analyses, separately for children with autism and controls, and separately for the raw scores, low-threshold items, and high-threshold items of the Sensory Profile. In each case we examined the ability of the sensory profile scores to explain the sleep disturbance scores. Initial regression models contained all 5 predictors (i.e., one predictor from each sensory modality) together. These analyses were followed-up by regression models containing each of the predictors separately. This enabled us to determine the contribution of scores from each sensory modality. In a final regression analysis we also added age, gender, medication-use, and ADOS scores for the models to determine whether these variables had an effect of the variance explained.

## Results

### Sensory sensitivities in autism

Children with autism exhibited abnormal sensory sensitivities that were evident in the Sensory Profile scores of all five sensory modalities (Figure 1 & Table 2). The total raw scores from the Sensory Profile were significantly smaller in children with autism as compared to the control children in the auditory, visual, vestibular, touch, and oral domains (t(129) < −5.78, p < 0.001). Similar results were also evident when comparing only male children in the two groups (t(95) < −4.7, p < 0.001), when including only children with autism who completed an ADOS assessment (t(109) < −4.1, p<0.001), or when excluding children with autism who were taking medications (t(107) < −5.5, p < 0.001).

**Table 2:**
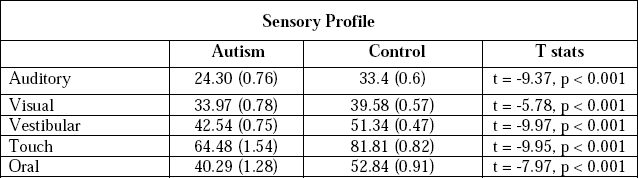
Sensory Profile scores. The mean and standard error (in parentheses) are presented for the autism (left column) and control (middle column) groups along with the statistics of two-sample t-tests with unequal variances (right column). P-values are Bonferroni corrected for multiple comparisons.

No significant differences were found in sensory sensitivity scores between children with autism who were or were not taking medication (t(41) < 0.9, p > 0.34), nor between male and female children with autism (t(20) > −1, p > 0.29).

**Figure 1:**
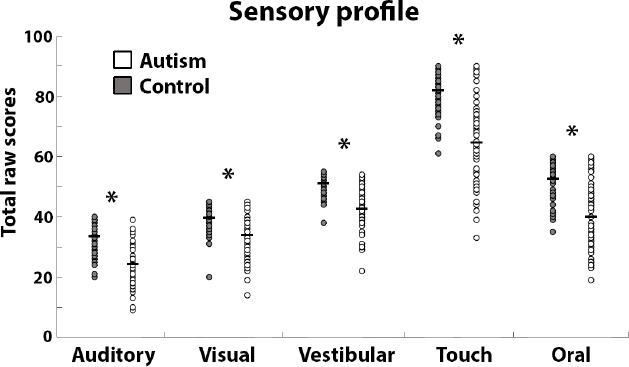
Scatter plots of sensory profile scores. The total raw scores from each sensory modality in the Sensory Profile are presented for children with autism (white) and controls (gray). Each circle represents a single child. Black lines: group mean. Asterisks: significant differences across groups (p < 0.001, Bonferroni Corrected).

The total raw scores of the Sensory Profile are the sum of scores reported for both low and high threshold items on the questionnaire, which measure hyper and hypo sensitivities respectively (Fig. 2). Note that children can be hyper-sensitive to some stimuli and hypo-sensitive to other stimuli even within the same sensory domain. Children with autism exhibited significantly lower scores on both the low (t(129) < −4.55, p < 0.005) and high (t(129) < −5.75, p < 0.005) threshold items in all five sensory modalities.

Performing the same analyses only with males revealed equivalent results for both low (t(95) < −4.1, p < 0.001) and high threshold items (t(95) < −4.6, p < 0.005). Similar results were found when performing the analysis only with children who had ADOS scores for both low (t(109)<−3.65, p<0.01) and high threshold items (t(109)<−4.5, p<0.001). Excluding children who were taking medication from the autism group also revealed equivalent results for both low (t(107) < −4.8, p < 0.001) and high threshold items (t(107) < −5.2, p < 0.001).

**Figure 2:**
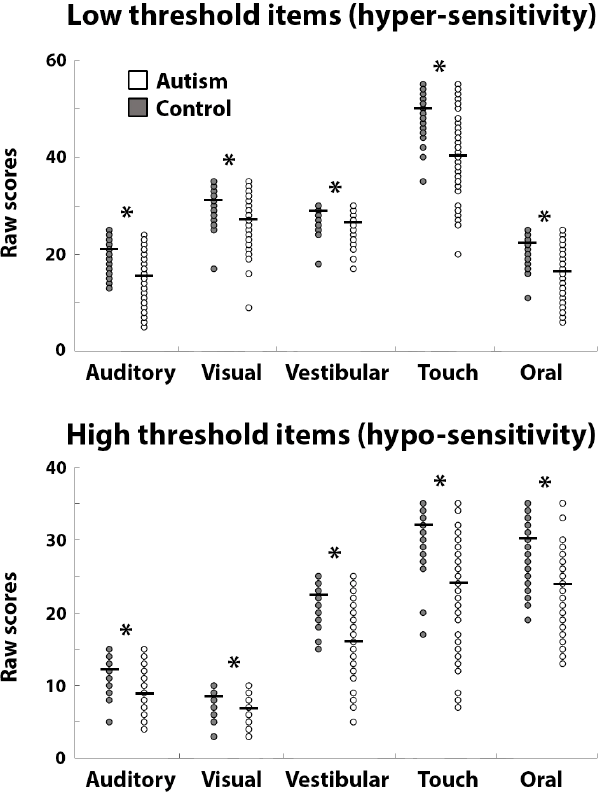
Hypo and Hyper sensitivities. Scatter plots of low (top) and high (bottom) threshold item scores from the Sensory Profile for children with autism (white) and controls (gray) groups. Each circle represents a single child. Black lines: group mean. Asterisks: significant differences across groups (p < 0.005, Bonferroni Corrected).

Both autism and control groups (Table 3) exhibited strong and significant correlations across Sensory Profile scores in the different sensory domains, demonstrating that sensory sensitivities of individual children were similar across sensory domains in both groups.

**Table 3:**
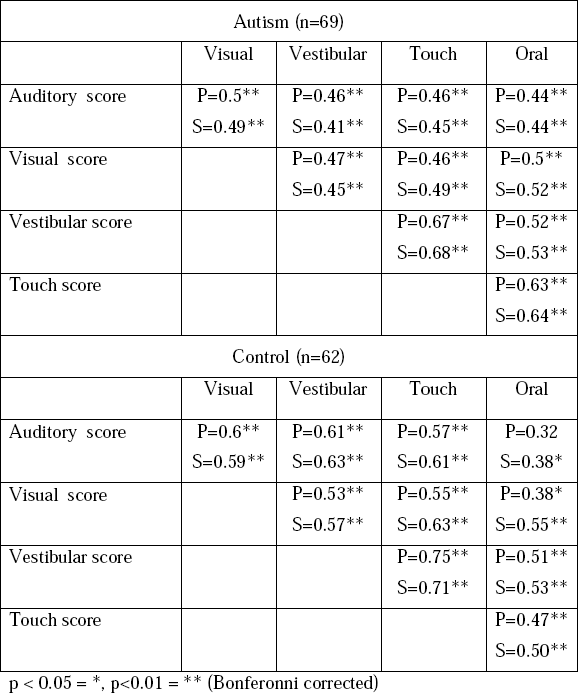
Correlations across sensory domains. Pearson’s (P) and Spearman’s (S) correlation coefficients are presented for each pair of sensory domains in the Sensory Profile for the autism (top) and control (bottom) groups.

### Sleep problems in autism

Children with autism exhibited significantly larger sleep disturbance scores than control children in the bedtimes resistance (t(129) = 5.35, p < 0.001) sleep onset delay (t(129) = 6.02, p < 0.001), sleep duration (t(129) = 4.88, p < 0.001), sleep anxiety (t(129) = 4.54, p < 0.001), night wakings (t(129) = 3.27, p<0.001), and parasomnia (t(129/) = 5.13, p < 0.001) subscales (Table 4 & Figure 3). Total sleep disturbances were also significantly larger in children with autism as compared to control children (t(129) = 5.86, p < 0.001, Table 4 & Figure 3). The scores in the other CSHQ subscales (sleep disordered breathing, and daytime sleepiness) were not significantly different across groups. A total sleep disturbances score of 41 is considered to be a useful clinical cutoff when screening children for sleep problems [27]. Significantly more children with autism (85.5%) had scores higher than this cutoff in comparison to control children (54.8%) (X^2^(1) = 14.91, p<0.001).

**Table 4:**
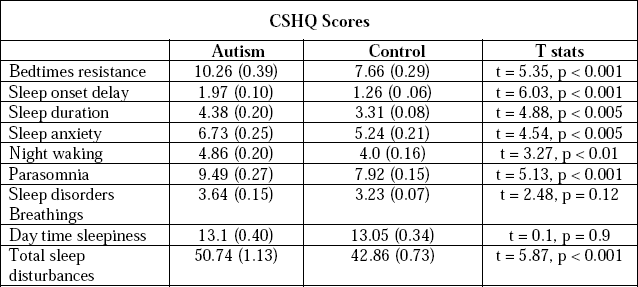
CSHQ scores. The mean and standard error (in parentheses) are presented for the autism (left column) and control (middle column) groups along with the statistics of two-sample t-tests with unequal variances (right column). P-values are Bonferroni corrected for multiple comparisons.

Performing the same analysis while including different subsets of children with autism yielded equivalent results. When excluding children with autism who were taking medication, there were significantly larger scores in the autism group in bedtimes resistance (t(107) = 2.124, p < 0.05) sleep onset delay (t(107) = 2.41, p < 0.05), sleep duration (t(107) = 2.9, p < 0.01), and parasomnia (t(107) = 2.59, p < 0.05) subscales as well as total sleep disturbances (t(107) = 3.23, p < 0.01). When including only male children in both groups, there were significantly larger scores in the autism group in bedtime resistance (t(95) = 4.24, p < 0.001), sleep onset delay (t(95)=4.31, p< 0.001), sleep duration (t(95) = 4.23, p < 0.001), sleep anxiety (t(95)=3.17, p<0.01) and parasomnia (t(95 = 4.21, p < 0.001) subscales as well as total sleep disturbances (t(95) = 5.09, p < 0.001). When including only children who completed the ADOS in the autism group, there were significantly larger scores in the autism group in bedtime resistance (t(109) = 4.14, p < 0.001), sleep onset delay (t(109)=5.05, p< 0.001), sleep duration (t(109) = 3.68, p < 0.001), sleep anxiety (t(109)=3.43, p<0.01) and parasomnia (t(109) = 4.28, p < 0.001) subscales as well as total sleep disturbances (t(109) = 4.3, p < 0.001). In this final analysis the night waking scores were almost significantly larger in the autism group (t(109) = 2.7, p = 0.06).

**Figure 3:**
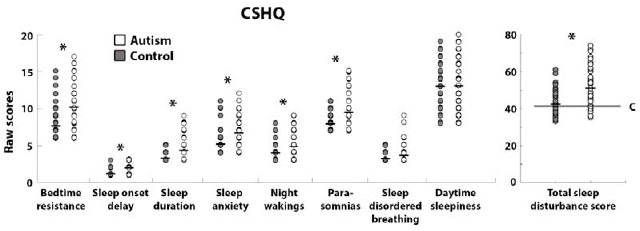
Scatter plots of CSHQ scores. Specific subscales (left) and total sleep disturbances (right) are presented for children with autism (white) and controls (gray). Each circle represents a single child. Black lines: group mean. Asterisks: significant differences across groups (p < 0.01, Bonferroni Corrected). C: cutoff line that is often used when screening children for clinical sleep problems [27].

### The relationship between sleep disturbances and sensory sensitivities

We examined the relationship between the total sleep disturbances score and sensory sensitivity scores in each of the five sensory systems (Figure 4, left panel). Significant negative correlations were apparent between the touch or oral sensitivity scores and total sleep disturbances scores of children with autism when computing Pearson’s correlations (r = −0.55, p < 0.001 and r = −0.42, p < 0.005) or Spearman’s correlations (r = −0.53, p < 0.001 and r = −0.41, p < 0.005). Control children exhibited significant negative correlations between vestibular or touch sensitivity scores and total sleep disturbance scores when computing Pearson’s correlations (r = −0.45, p < 0.005 and r = −0.42, p < 0.01). Spearman’s correlations exhibited similar trends, but were not significant (r = −0.31, p < 0.07 and r = −0.29, p < 0.1). All other correlations were not statistically significant.

**Figure 4:**
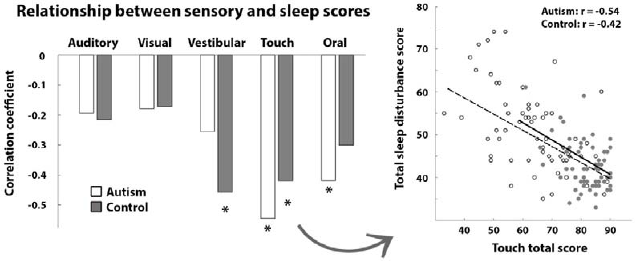
Relationship between Sensory Profile and CSHQ scores. Left panel: Pearson correlations between sensory scores from each of the five sensory modalities and total sleep disturbance scores in the autism (white) and control (gray) groups. Right panel: Scatter plot of the touch versus sleep scores. Each circle represents a single child. Asterisks: significant correlation (p < 0.05, Bonferroni corrected)

Despite the correlation across sensory subscale scores (Table 3), only ***some*** of the sensory scores were significantly correlated with the total sleep disturbance scores (Figure 4 & Table 5). Furthermore, while significant negative correlations were apparent in the touch domain of both groups, scores of children with autism were distributed over a much wider range of values than scores of control children (Fig. 4, right panel). This means that the strong correlations in the autism group represented a tight relationship that was apparent also in cases of severe sleep disturbances and sensory problems (i.e., correlation was robust throughout the larger range).

In additional analyses we examined whether the sleep disturbance scores were more strongly associated with low or high threshold items from the Sensory Profile (Figure 5,Table 5). Significant negative correlations were apparent between low item scores (i.e., hyper sensitivity) in the touch and oral domains of children with autism (r = −0.5, p < 0.001 and r = −0.35, p < 0.02 respectively) and in vestibular and touch domains of controls (r = −0.46, p < 0.001 and r = −0.42, p < 0.01). Low item scores in all other sensory domains were not significantly correlated with sleep disturbance scores. Significant negative correlations were also apparent between scores of high items (hypo sensitivity) in the touch and oral domains of children with autism (r = −0.41, p < 0.005 and r = −0.42, p < 0.005). High item scores in all other sensory domains of children with autism and all sensory domains in control children were not significantly correlated with sleep scores.

**Figure 5:**
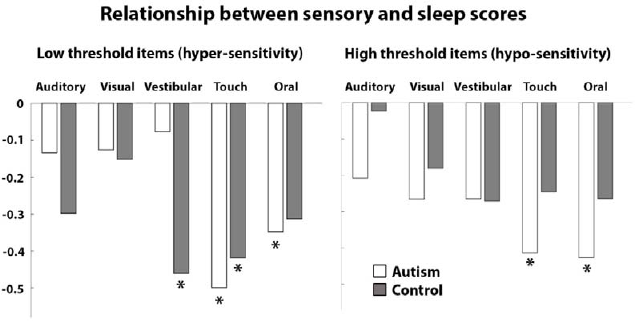
Relationship between total sleep disturbance scores and hypo or hyper sensitivity in each of the sensory domains. Low-threshold (left) and high-threshold (right) items that measure hyper and hypo sensitivity respectively in the autism (white) and control (gray) groups. Asterisks: Significant correlations (p < 0.05, Bonferroni corrected).

**Table 5:**
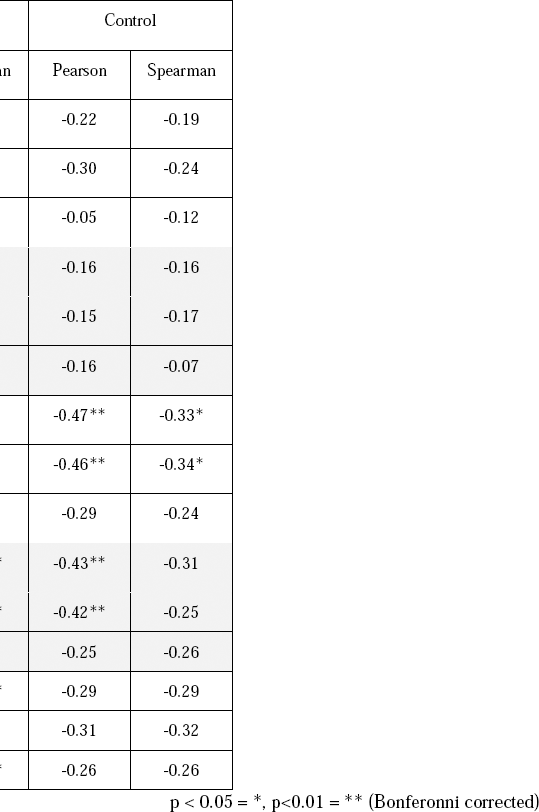
Relationship between total sleep disturbance scores and total, hypo, or hyper sensitivity in each of the sensory domains. Pearson’s and Spearman’s correlations between the total, high-item, and low-item scores of the Sensory Profile and the total sleep disturbance score of the CSHQ for children with autism (left) and controls (right).

### Explaining sleep disturbances with sensory sensitivity scores

In a final set of regression analyses, we quantified how much of the variance in sleep disturbance scores could be explained by the total raw scores from each of the five sensory domains separately and also when including all of them together. Incorporating the total raw scores from all of the sensory domains into a single regression model yielded an adjusted R^2^ value of 0.293 in the autism group and 0.198 in the control group (Table 6).

When performing the regression with a single predictor from each sensory modality separately, the touch scores stood out in their ability to explain sleep disturbance scores in children with autism (adjusted R^2^ = 0.286). This suggests that the touch scores could single-handedly explain as much of the variance in sleep scores as the integrated model, which contained all 5 predictors. In the control group, the vestibular and touch scores yielded adjusted R^2^ of 0.196 and 0.162 respectively, thereby demonstrating their ability to explain large portions of the variability in sleep scores in the control group.

We performed equivalent analyses while separating the scores from the low-threshold (hyper sensitivity) and high-threshold (hypo sensitivity) items. The full low-threshold regression model (containing 5 predictors/modalities) yielded adjusted R^2^ values of 0.243 and 0.206 in the autism and control groups respectively while the full high-threshold model yielded adjusted R^2^ values of 0.198 and 0.074. This demonstrates that the ability of Sensory Profile scores to explain sleep disturbance scores comes from low-threshold items/questions that measure sensory hyper sensitivity.

When examining low-threshold items in each sensory modality separately, the touch domain again stood out in its ability to explain sleep disturbance scores in children with autism (adjusted R^2^ = 0.238). This result demonstrates that low-threshold touch scores could single-handedly explain most of the variance that was explained by the full model with the total raw scores from all sensory modalities.

Adding age, gender, medication usage, and ADOS scores as additional predictors to the full models that contained all five predictors, had negligible effects on the results. The adjusted R^2^ value improved from 0.293 to 0.295, demonstrating that these additional predictors explained only 0.2% of the variance in the sleep disturbance scores.

Taken together, these results suggest that total sleep disturbances in children with autism are most strongly associated with hyper sensitivity towards touch. In contrast, sleep disturbances in control children are most strongly association with vestibular hyper sensitivity. Notably, scores in the visual and auditory sensory domains were remarkably weak in explaining sleep disturbances in both groups.

**Table 6:**
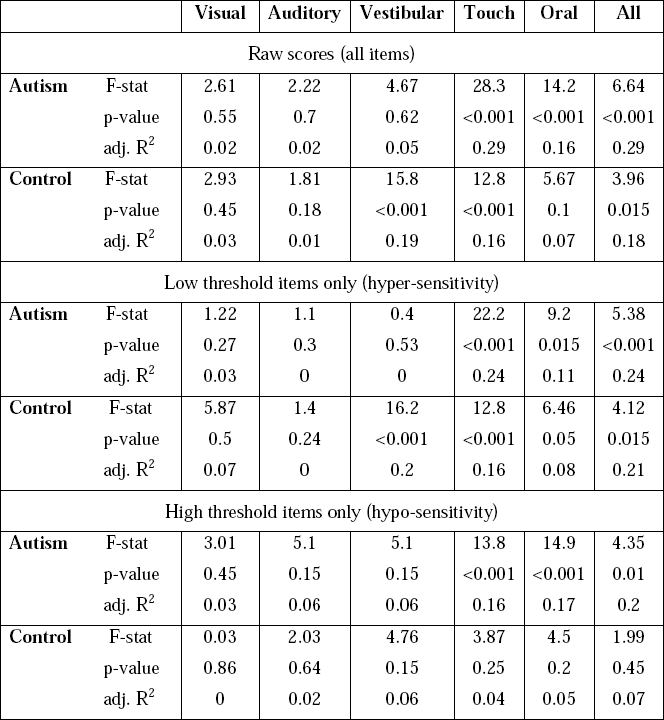
Variance explained by Sensory Profile scores. Results of regression analyses using six different models: one model with a single predictor for each of the sensory modalities and a sixth model containing all 5 predictors together. This analysis was performed once with the total raw scores (i.e., sum of low and high threshold items/scores) and again with the low and high threshold items separately. F statistics, p-values (Bonferroni corrected), and adjusted R^2^ are presented for each model.

## Discussion

Our results reveal a strong, significant, and specific relationship between sleep disturbances and touch and oral sensory abnormalities in autism (Figures 4-5). In particular, hyper sensitivity towards touch exhibited the strongest relationship, by single-handedly explaining 24% of the variability in sleep disturbance scores (Table 5). Similar relationships were also evident in the control group where sleep disturbances were strongly associated with touch and vestibular hyper-sensitivities (Figure 5). While one cannot infer causality from correlations, we speculate that hyper sensitivity towards touch may interfere with sleep onset and sleep maintenance in children with autism, thereby generating severe sleep disturbances in children with severe hyper sensitivities towards touch. Further studies examining the potential benefits of tactile sensory therapies/aids on sleep quality and associated behavioral improvements could potentially establish such causality and improve clinical care for children with autism who exhibit these particular symptoms.

### Specificity of sensory abnormalities associated with sleep disturbances

Are sleep disturbances associated with a general multi-modal sensory problem in autism, or with hyper sensitivity in a particular sensory domain? Recent studies using the short Sensory Profile have suggested that sleep disturbances are weakly associated with general sensory abnormalities, which explain 1-8% of the variance in sleep disturbance scores [19,23]. The short Sensory Profile, however, does not allow one to separate sensory hypo and hyper sensitivity scores nor scores of individual sensory modalities.

A more in-depth assessment using the complete Sensory Profile, reveals that sleep disturbances are not equally associated with sensitivities in all sensory modalities. In contrast to the strong relationship between sleep disturbances and abnormalities in the touch domain, sensory abnormalities in the visual and auditory domains were not associated with sleep problems in either the autism or control groups (Figures 4-5). Furthermore, sleep disturbances were more strongly associated with hyper-sensitivities towards touch than hypo-sensitivities (Table 4). Our results, therefore, clearly demonstrate that sleep disturbances are associated with sensory abnormalities in specific sensory modalities and cannot be generalized across all sensory domains.

### The relationship between sensory problems and sleep disturbances in typical development

Our findings are in line with several previous studies, which have also reported significant relationships between sensory hyper sensitivity on the Sensory Profile and sleep disturbances in infants [34] children [35] and adults [36] with typical development. Two of these studies examined sensitivity scores separately in each of the sensory modalities and reported that tactile hypersensitivity scores explained the largest amount of variability in sleep disturbance scores (~25%), while scores in other sensory domains such as vision and audition, explained a considerably smaller portion of the variability [35,36]. In our study both vestibular hyper sensitivity and hyper sensitivity toward touch scores could single handedly explain a considerable amount of the variability in sleep disturbance scores of control children (20% and 16% respectively). Taken together, accumulating evidence suggests that hyper sensitivity in the tactile and vestibular modalities are particularly useful for explaining sleep disturbances in children with typical development.

### Conclusions and future directions

The findings reported in this study resonate well with the hyper arousal theory of insomnia. The theory suggests that increased levels of anxiety/stress lead to cortical over-reactivity to sensory stimuli, which interferes sleep onset and sleep maintenance [37]. Evidence for this theory comes, in part, from studies that report abnormally strong responses to sensory stimuli right before sleep onset and during different stages of sleep as measured by EEG responses to sounds [38–40]. Interestingly, an analogous theory suggests that autism may be caused by the abnormal development of hyper-aroused and over-responsive neural circuits [41,42].

Since parental questionnaires may be biased by parental sensitivities, there is strong motivation to conduct further studies that will examine whether abnormally large tactile responses (as measured by psychometric and neuroimaging techniques) are apparent during wake and sleep periods in children with autism who exhibit sleep disturbances. Such research could further characterize individual children with autism who require specific interventions for treating tactile hyper sensitivity.

Dividing the heterogeneity of autism and identifying individuals with shared symptoms and etiologies is a major goal of contemporary autism research [2,43,44]. With this in mind, we demonstrate that sleep abnormalities are specifically tied to touch hyper-sensitivities in young children with autism. This combination of characteristics identifies specific children with autism who may benefit from specific tactile interventions/aids.

## Abbreviations

CSHQ: Child Sleep Habit Questionnaire
ATN: Autism Treatment Network
ADOS: Autism Diagnostic Observation Schedule

